# The self-renewal procedures of mesenchymal stem cells in the blood

**DOI:** 10.1101/2023.02.03.527029

**Authors:** Wuyi Kong, Hong Wang, XiaoPing Zhu, XiuJuan Han

## Abstract

**Background:** Although mesenchymal stem cells (MSCs) are most commonly used in cell therapy and stem cell research, the mechanism and the locations of their self-renewal are still unknown.

**Method:** Mouse blood was collected, and examined under microscopy. The results were compared with the data of human umbilical cord blood (hUCB) collected 10 years ago.

**Results:** We found that the procedure of self-renewal for the mesenchymal stem cells in mouse blood and hUCB needs at least 5 steps. First, tube-shaped stem cell niches release long segmented materials composed of sand-like particles and semitransparent granules. Second, the sand-like particles and semitransparent granules separate from the segmented materials. Third, each of the individual semitransparent granules releases groups of fusiform-shaped structures that do not stain to H&E. The sizes of the fusiform-shaped structures range from 1 to 100 μm in length in mouse blood, but can be 200 μm in hUCB. Fourth, the large-sized fusiform structures can directly transform into lineage-restricted cellular structures; the medium-sized fusiform structures fuse or engulf each other to form cellular structures. The cellular structures further acquire membranes from the adjacent nucleated cells. Fifth, the nucleolus appears in the new cellular structures before the nucleus. During all the procedures, the adjacent nucleated mesenchymal cells are must needed. Thus, these newly formed cellular structures will further differentiate into nucleated mesenchymal stem cells.

**Conclusion:** Our findings again provide new evidence that, in physiological conditions, mesenchymal stem cell self-renewal needs several steps to complete, which, however, does not occur by mitotic division. The tube-shaped structures are the niches of the stem cells.

## Background

For decades, mesenchymal stem cells (MSCs) have been the most commonly used “words” for the cells in cell therapy and stem cell research. Although they have been described to have multilineage functions to regenerate osteoblasts, adipocytes, chondrocytes, myocytes, and even neuroglia (George et al., 2019; Petersen et al., 1999), the opposing openings that question their multilineage function have never been stopped because the identification markers for MSCs are not specific (Dominici et al., 2006; Lin et al., 2013). Additionally, more evidence indicates that different tissues have their own tissue-specific stem cells to meet their requirements (Rinkevich et al., 2015; Visvader and Clevers, 2016); thus, MSCs isolated from different tissues may not be the same. In addition, a recent report claimed that restricted lineage segregation occurs during early embryonic development (Wuidart et al., 2018), indicating that the cell lineages have been carefully categorized prenatally or that cells switch from multipotent to unipotent in embryonic development. Thus, it is important to elucidate the potency of postnatal MSCs.

New cells are constantly needed in the body to replace aged cells and repair wounds, which suggests that all types of cells require self-renewal to meet life-long requirements. Thus, elucidating where and how the cells are renewed have become the very important research topics. The concept that a microenvironment called the niche is the place where the stem cells are either located or renewed has been well accepted. Most reports have described stem cell niches located in bone marrow sinusoidal, perivascular or specific organs (Morrison and Scadden, 2014). These niches are composed of different types of cells (Ding et al., 2012) or composed of cells and extracellular structures such as the extracellular matrix (ECM) (Adams et al., 2006; Malara et al., 2014). Until now, the complexities and structures of niches have remained largely unknown. How stem cells are renewed is also unclear. Some reports have shown that stem cells in niches are in a quiescent state that can be activated by various factors or hormones and undergo a selfrenewal pattern called asymmetric division to maintain life-long production of mature cells (Berika et al., 2014; Fuchs and Chen, 2013; Li and Clevers, 2010; Moore and Lemischka, 2006). Apposing report utilizing imaging techniques in live mice indicated that dermal stem cells do not undergo asymmetric cell division during differentiation (Rompolas et al., 2016). Thus, the mechanism of postnatal stem cell self-renewal is still unclear because the presented statements are mostly displayed by schematic diagrams.

In our previous reports, we found two tube-shaped structures. One of them releases a bundle of thin filament-like structures, or “progenitor-releasing filaments”, and another releases a group of bud-like structures (Kong et al., 2019). Each of them can release dozens of morphologically identical spore-like progenitors, one of which further becomes CD34-positive cells. Because each of these structures produces and releases only one distinct spore-like cellular type or progenitors, we believe that the lineage of these progenitors has been predetermined when their tube-shaped structures or niches are built. Also, in our previous report of cultured hUCB (Kong et al., 2013a), we observed many small particles and similar fusiform structures that we termed “Non-platelet RNA-containing particles (NPRCPs) and “particle fusion-derived non-nucleated cells (PFDNCs), respectively. The small particles could fuse with each other to form small cellular structures, and two or more particle-derived small cells could further fuse or engulf each other to become larger-sized Oct4- and Sox-2-positive pre-stem cells. However, in our previous study, we did not know the origins of these small particles.

Here, we present the renewal procedures of the blood mesenchymal cells in physiological conditions. We found that the production of mesenchymal cells occurs inside one type of tightly sealed tube-shaped structures or niches that are lineage predetermined. Only one type of cell was developed from a tube-shaped structure. Thus, blood mesenchymal cells are unipotent cells. The renewal of these cells is not due to any type of mitotic division, whether symmetric or asymmetric.

## Methods

### Animals and materials

Male Balb/C and C57BL6 mice at 8 to 16 weeks of age were purchased from Vital River Laboratories (Beijing). Mice received food and water *ad libitum*. The animal committee in Khasar Medical Technology approved the research protocol “Stem cells in the mouse blood”. All animal procedures followed the guidelines of the Animal Regulatory Office relevant to national and international guidelines.

### Mouse blood collection

Five of 8- to 10-week-old mice of each strain were used for each tissue preparation. Blood was collected from each mouse via cardiac puncture under sterile conditions after CO2 euthanasia. Each syringe was prefilled with 100 μl of heparin (0.5%) to prevent coagulation. Blood was placed on ice for ~5 min, pooled (~5 ml), transferred to a 50-ml centrifuge tube, immediately diluted with 1x phosphate buffered saline (PBS) at a ratio of 1:5 (blood:PBS), and then centrifuged at 200xg for 5 min. After removing the supernatant, the bottom cellular portion was diluted with PBS in a 1:1 ratio, and then, fixed at 4°C overnight after slowly adding 15% paraformaldehyde to a final concentration of 4% paraformaldehyde. The next day, the cells were centrifuged at 200xg for 5 min, washed 2 times with PBS and centrifuged at 100xg for 5 min and twice at 50xg for 5 min to remove free erythrocytes. Visible red clots were removed during washing. The cellular portions of each centrifugation were pooled and examined. After a final centrifugation and removal of the supernatant, the cellular portion was fixed again with 4% paraformaldehyde at a ratio of 1:5 (original blood volume:fixative volume) and dropped (200 μm/drop from ~5 cm above) onto gelatin-coated histology slides. At least 20 slides from each preparation of each strain of mice were stained and examined.

### Hematoxylin and eosin (H&E) staining

Cellular portions from each mouse strain preparation were dropped on at least 20 slides. Slides were dried overnight, washed 3 times with PBS, and stained with H&E. Slides were visualized by conventional fluorescence microscopy (Leica, Wetzlar, Germany) and photographed with the use of a digital camera (Leica, DFC500).

### Human umbilical cord blood (hUCB)

No new hUCB was collected for the current study. Slides stained for our previous hUCB study (Kong et al., 2013a) were reexamined. Digital movies for the cultured hUCB cells were taken 10 years ago in the same study (Kong et al., 2013a). Briefly, ten ml hUCB were collected and centrifuged at 200 xg form 10 min. After removing plasma, blood was incubated in erythrocyte lysis buffer (155 mM NH4Cl, 10 mM KHCO3 and 0.1 mM EDTA) for 20 min, then underwent centrifugation at 300 xg for 10 min; the supernatant was transferred to a fresh bottle and centrifuged at 1000 xg. After a washing with phosphate buffered saline (PBS), pellets were cultured on collagen-coated plates in a-MEM containing 20% fetal bovine serum and antibiotics in a 5% CO_2_ humid incubator at 37°C. Medium was changed every day for the first 3 days to remove erythrocyte membranes and dead cells and every 2 days thereafter. Images were taken on the cells from 1 - 3 weeks after cultural.

## Results

### Identification of segmented materials

Mouse blood was collected, prepared, stained, and carefully examined under a microscope. During the observations, we often observed many dark-colored segments that were initially thought to be the contaminations of the dyes or others during the blood preparation. However, after three filtrations for the dyes before each stain and careful performance of the sample preparation procedures, the dark-colored segments were still observed in every separated mouse blood preparations. Under microscopy, we found that the sizes of these segments, although were differ in length, but similar in width (S. Fig. 1), suggesting that they were separated from some long structures that had similar widths. One segment was more than 400 μm in length (Fig. 1A), incompact, and composed of dark sand-like particles and semitransparent larger granules. A cracked line appeared (arrows in Fig. 1A) in the middle of this segment, suggesting where a further breaking could occur. The shorter segments continued to disperse (arrows in Fig. 1B-1C) and eventually separated and exposed the individual semitransparent larger granules that were approximately 10 to 20 μm in diameter (Fig. 1D-1F).

**Figure 1.**
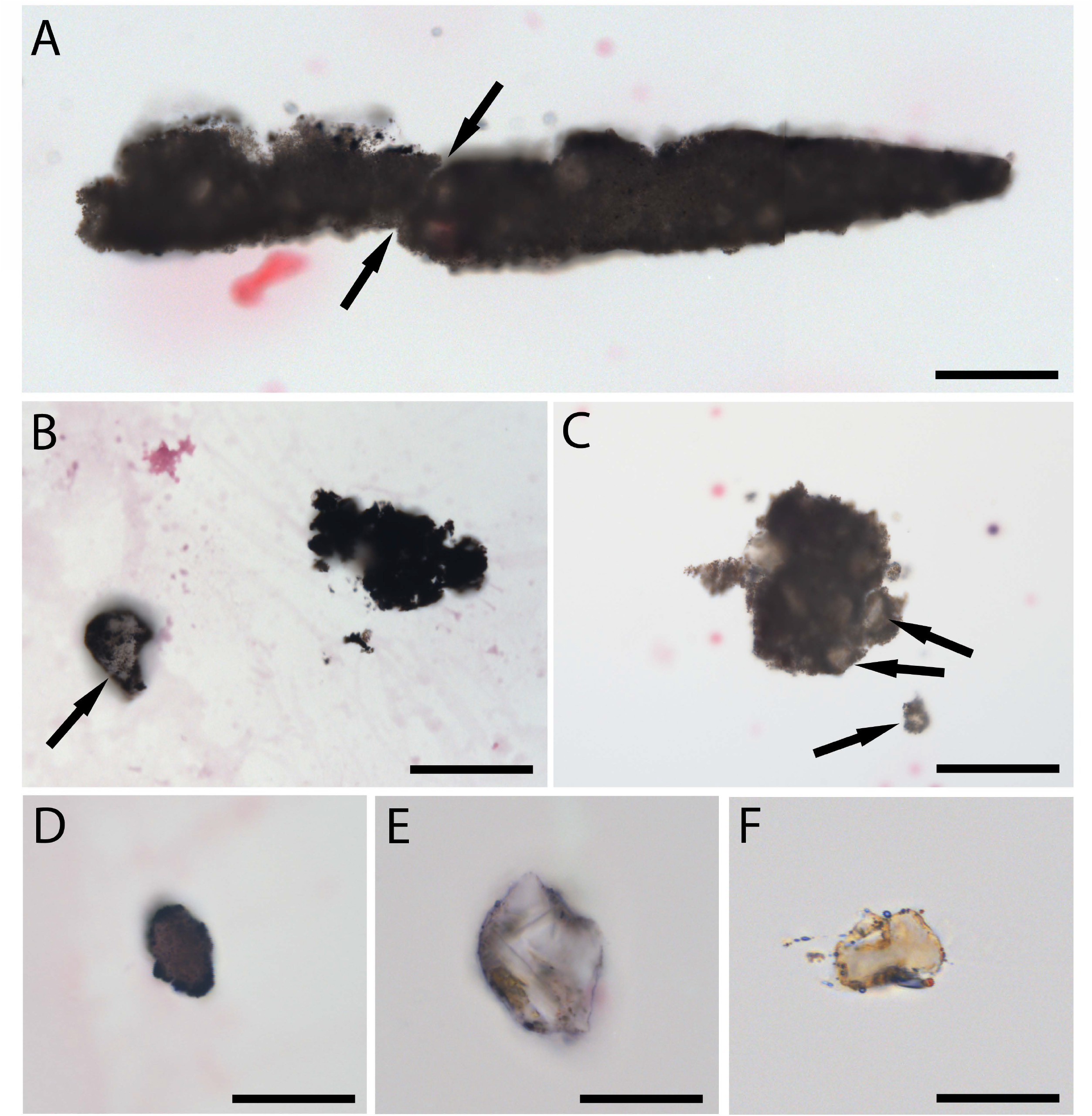
Dark segmented materials in mouse blood. Dark-colored segmented material that was approximately 400 μm in length (A) was observed in the blood sample. A cracked line appeared (arrows in A) in the middle of the long black material. The segmented materials contain semitransparent large granules (arrows in B – C) that were surrounded by almost black-colored materials. The semitransparent granules can be separated from the segments and sized approximately 15 to 20 μm in diameter (D – F). Bars in A-C = 50 μm; in D-F = 20 μm.

### Fusiform-shaped structures were released from semitransparent granules

We then observed numerous fusiform structures that were extruding from the dark-colored segments (Fig. 2A, 2B). The sizes of these fusiform structures were obviously different, which ranged from less than 1 μm to near 80 μm in length. Further, the images of the individual semitransparent larger granules showed clearly that each grouped fusiform structures were from one semitransparent larger granule (2C-2G). The outer membranes of these individual semitransparent granules were either in light-brown color (Fig. 2C, 2D) or transparent (Fig. 2E-2G), suggesting that the dark color was from the sand-like particles that had shed off from the surface of these semitransparent granules. We hypothesize that these dark sand-like particles may play important roles in providing nutrients to the semitransparent granules. Each grouped fusiform structure is arranged mainly in a line pattern, suggesting they were erupted out in one direction by force (Fig. 2A, 2H). To compare the size differences of these fusiform structures, all images in Fig. 2 were taken under the same magnification. The larger ones were approximately 80 μm in length (Fig. 2A), and the small ones were less than 1 μm (Fig. 2H). Due to the small sizes of these fusiform structures, they were mostly found floating on the top level of the tissue samples on tissue slides (Fig. 2H, S. Fig. 2). Additionally, they could not stain to H&E. These characteristics made most of them were hard to be identified under standard microscopic transmission light.

**Figure 2.**
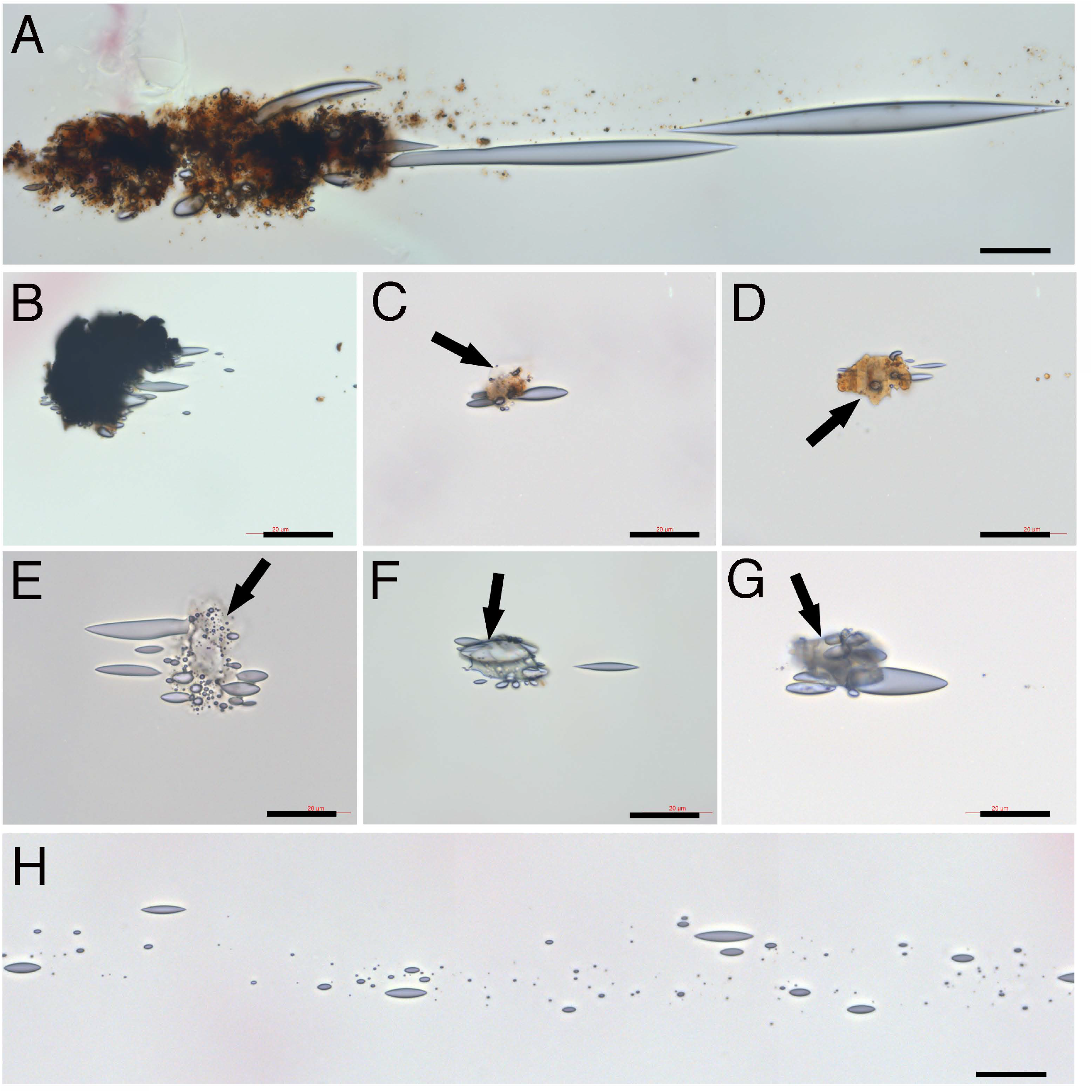
Fusiform-shaped structures were released from dark segmented materials. Numerous semitransparent fusiform-shaped structures were protruding from the black-segmented materials (A, B). Images of the individual semitransparent granules indicated that each group of fusiform structures was released from one semitransparent granule that had a broken membrane (arrows in C – G). The outer membranes of the semitransparent granules were either weak brown (C, D) or transparent (E – G). Many small fusiform structures were in line floating on the top level of the tissue samples on tissue slides (H). Bars = 20 μm.

### Wriggle-like contents appeared in the larger-sized fusiform structures

We observed that some larger fusiform structures showed light-contracted wriggle-like content inside (thick arrows in Fig. 3A-3C). The wriggle-like contents were mainly located in the center area of each fusiform structure. These contents were not observed in the images of the fusiform structures when they were released from the semitransparent larger granules (Fig. 2), but were observed in the separated larger-sized fusiform structures, suggesting the further development of fusiform structures after their release from the granules. Further, we observed that each large fusiform structure was connected to a round thin membrane that contained sand-like particles (thin arrow in Fig. 3C, S. Fig. 3), suggesting that the round thin membrane was related to the large fusiform structure. In order to elucidate their relations, different sources of microscopic light were applied. Due to not staining to H&E, the fusiform structures were difficult to be observed by bright transmission lights. Images were taken using either transmission light (Fig. 3D) or reflection light (Fig. 3E) on one fusiform structure-connected cellular structure. The round thin membrane was almost invisible by transmission light (Fig. 3D) but was observed by the reflection light, which showed that the end of the fusiform structure was located in the center of the round thin membrane. Thus, we hypothesize that these round thin membranes are the membranes of the semitransparent granules (Fig. 2) that degrade into small pieces after the release of the fusiform structures. The membranes of the semitransparent granules were not the standard cellular membrane, and could not stain to H&E. In comparison, two nucleated cells nearby stained positive to H&E.

**Figure 3.**
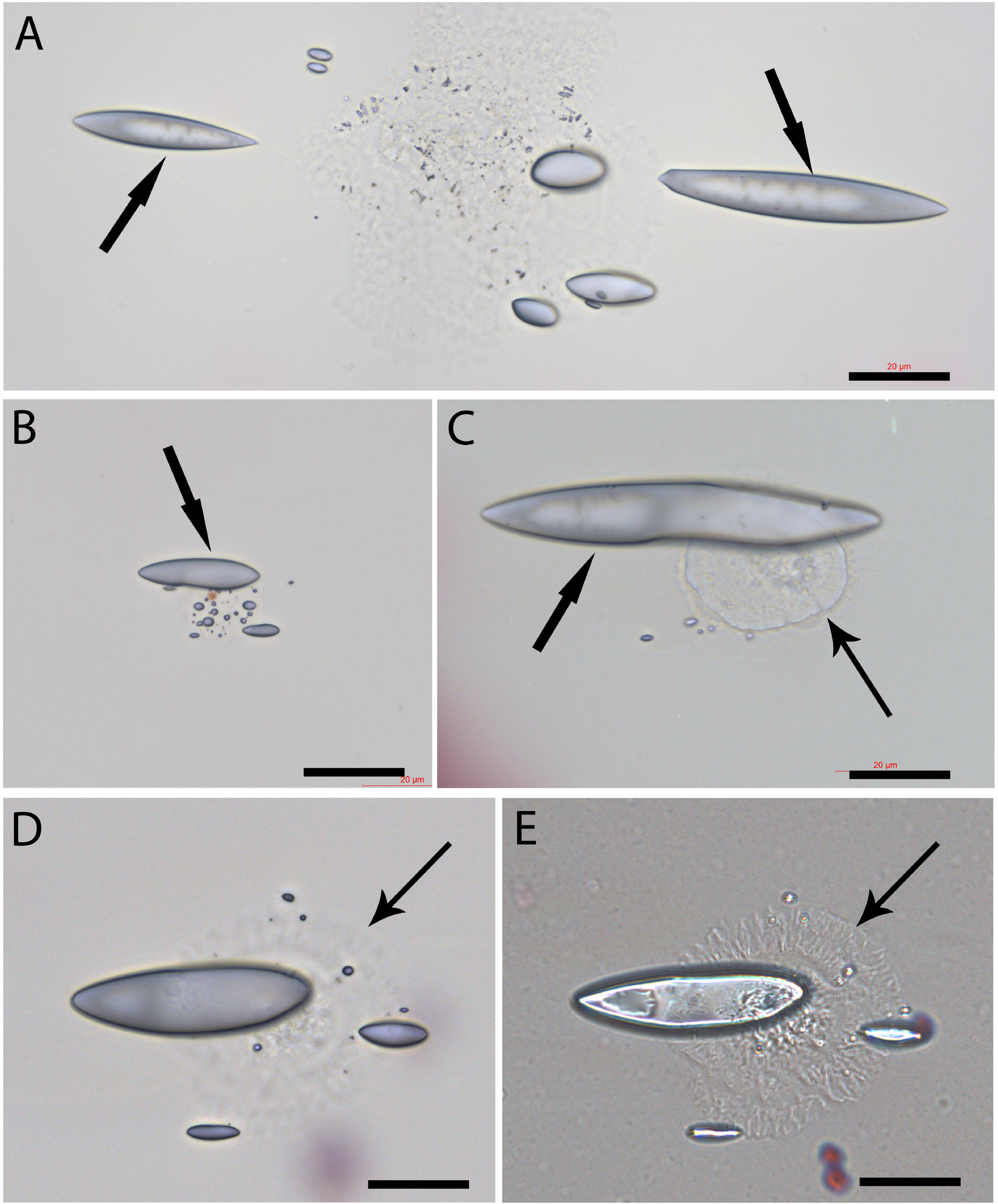
Wriggle-like contents appeared in the larger-sized fusiform structures. The sizes of the fusiform-shaped structures from one semitransparent granule were different. The larger fusiform-shaped structures showed wriggle-like contents inside (thick arrows in AC). A thin cellular structure is located right underneath a large content-containing fusiform structure (thin arrow in C). Images were taken using either transmission light (D) or reflection light (E) on the same cellular structure that was underneath a content-containing fusiform structure. Bars = 20 μm.

### Fusiform structures transformed into round cellular shapes

Using reflection light, we searched the tissue slides and found a large fusiform structure that was approximately 100 μm in length (B arrow in Fig. 4A). One line of cellular structures was located near it (C-E arrows in Fig. 4A), suggesting that they were related. High magnification images showed that some contents were at the end of this large fusiform structure (arrow in Fig. 4B). The cellular structures, although sized from small (arrows in Fig. 4C) to large (Fig. 4D-4E), were arranged in a line pattern. Except for the size differences, the morphologies of these cellular structures were similar, suggesting that they were the same lineage of cell progenitors. These cellular structures were not stained to H&E and did not show nuclei or standard cellular membranes, which made them undetectable by any dyes or surface markers. We believe that these cellular structures were transformed from a group of fusiform structures that had different sizes. The high magnification images showed numerous small particles located in the center of the large cellular structures (Fig. 4D-4F). The small particles were scattered in a well-arranged pattern around this cellular structure (arrows in Fig. 4F), which suggests that the small particles were popped out of the cellular structure during tissue preparations. These particles might be the sand-like particles surrounding the semitransparent larger granules and were taken by the fusiform structures after their release from the semitransparent granules. We assume that these small particles are also been called exosomes by some researchers and have attracted increased attentions in recent years in stem cell and therapeutic studies (Elahi et al., 2020).

**Figure 4.**
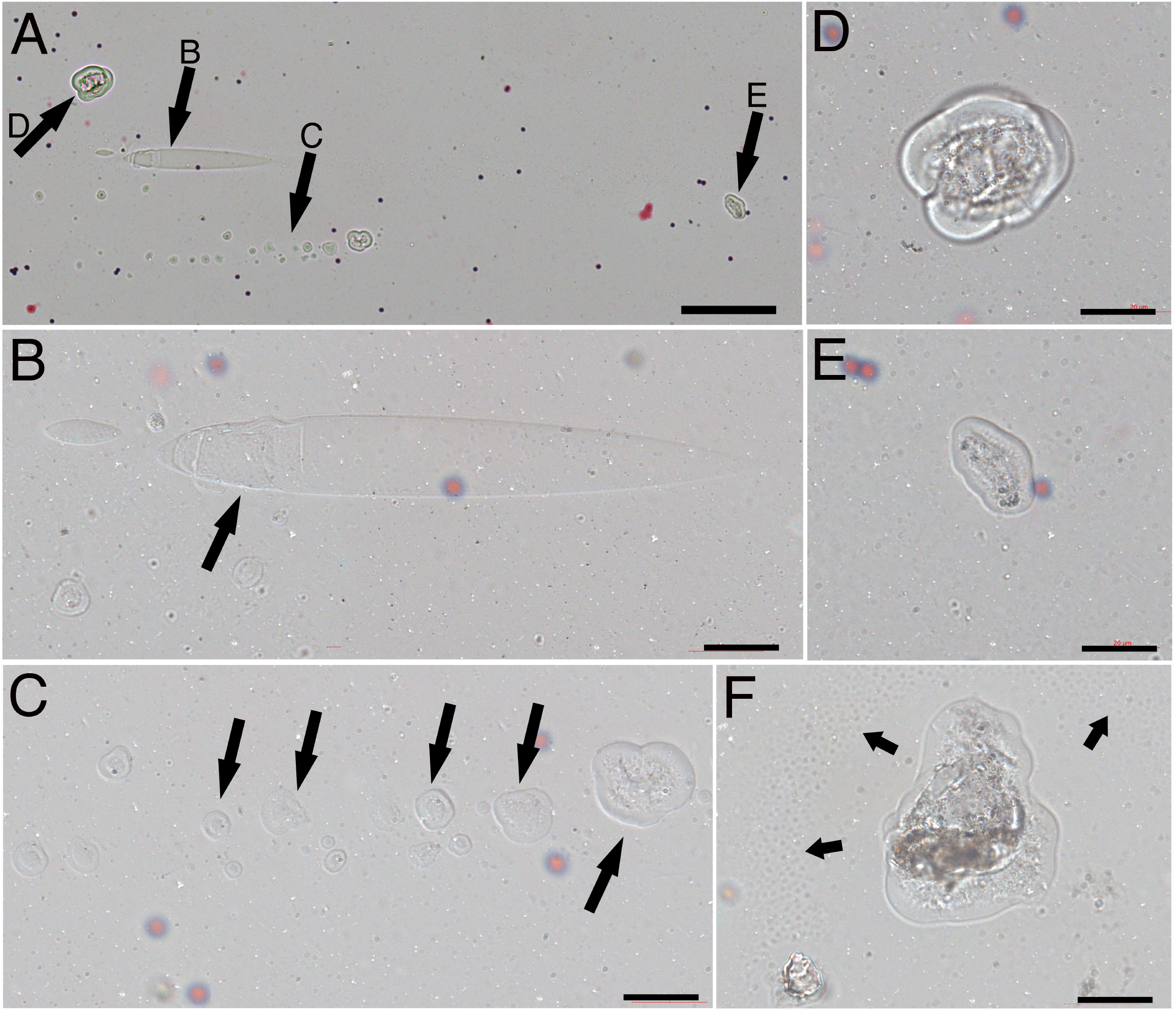
Observation of the cellular structures arranged in line. A large fusiform structure that was approximately 100 μm (arrow B in A) was identified. Many cellular structures were arranged in line near it (arrow C in A). Higher magnification images showed that the group of cellular structures sized differently and did not stain to H&E (arrows in C). The larger-sized cellular structures were found nearby (D-F). A large progenitor (F) that contained numerous small particles in the center area was surrounded by scattered small particles (arrows in F). Bar in A = 100 μm; in B – F = 20 μm.

### Fusiform structures in human umbilical cord blood (hUCB)

It has been well described that mesenchyme stem cells are rich in hUCB. Thus, mesenchyme stem cells used in many hUCB therapeutic studies were simply obtained by collecting the attached cells from the tissue cultural dishes. We then re-examined the tissue slides collected in our previous hUCB studies and found similar fusiform structures (Fig. 5). A group of fusiform structures closely located near a round thin membrane that contained numerous sand-like particles (Fig. 5A). A large fusiform structure and few small fusiform structures located next to a similar round thin membrane containing sand-like materials (Fig. 5B). Numerous small fusiform structures arranged in line were identified on top of the tissue sample slide (Fig. 5C). The shape and the arrangement of these fusiform structures in hUCB showed no differences from those in mouse blood, excepting that the size of large human fusiform structures (Fig. 5B) was longer than that in mouse blood. It was nearly 200 μm in length, while the longest one found in mouse blood was approximately 100 μm (Fig. 4B).

**Figure 5.**
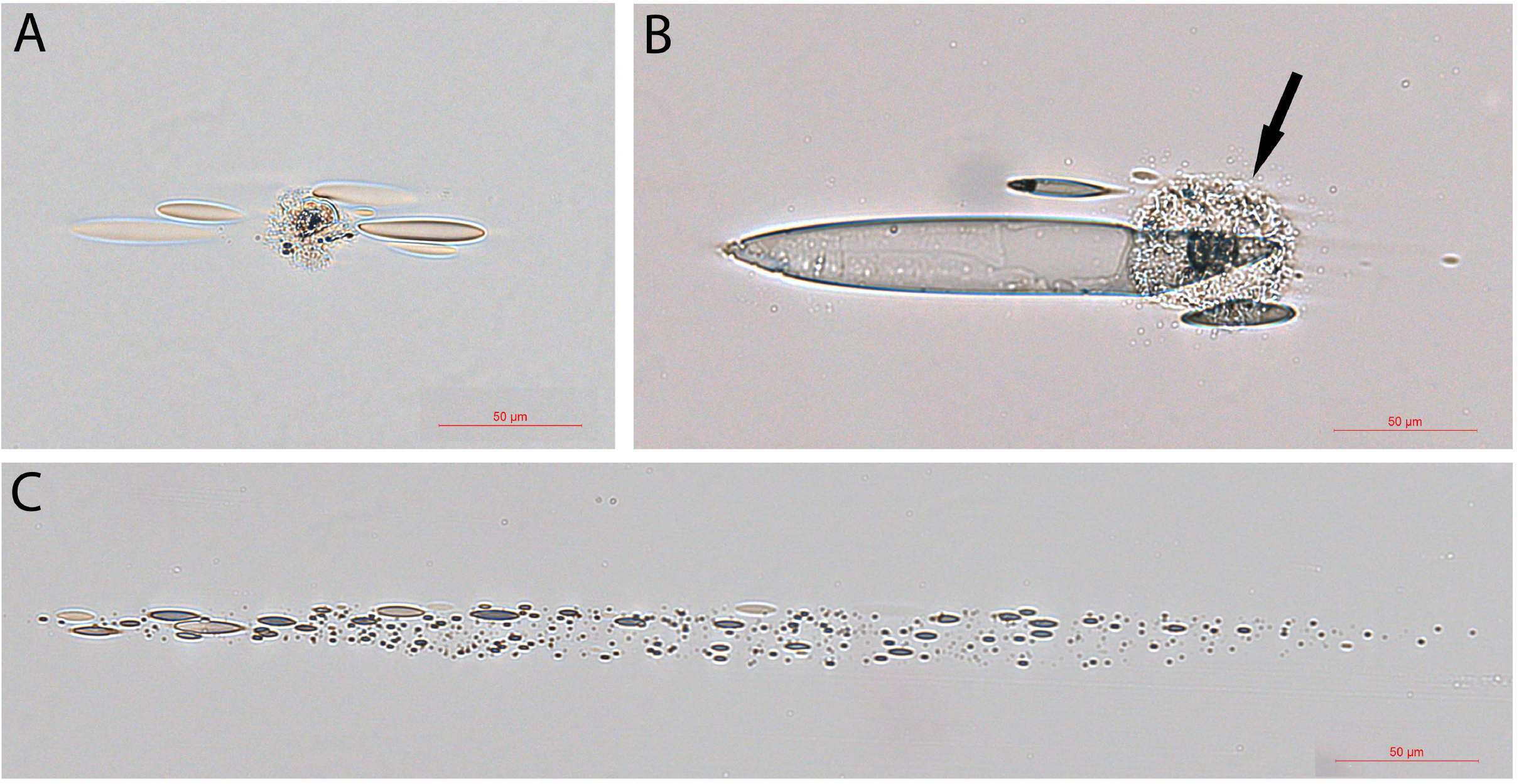
Fusiform structures were identified in human umbilical cord blood. In human umbilical cord blood slides, we identified groups of fusiform structures that were also connected to a thin membrane structure (A, arrow in B). The length of a large fusiform structure was about 200 μm (B). Many small fusiform structures arranged in line were identified on top of the tissue sample slide (C). Bars = 50 μm.

### Large fusiform structures directly transformed into mesenchymal cell progenitors

After identifying the fusiform structures in mouse blood and hUCB, we re-examined the video records from our previous cultured hUCB study and found that the large fusiform structures were directly connected to the cell-like structures (arrows in Fig. 6A, 6B, Movie 1), which suggest that the fusiform structures participated the formation of the cellular structures. The images clearly showed that the wriggle-like contents were in the center of each large fusiform structure. The middle panels showed the time-lapse images that a large fusiform structure (arrows) was connected to a cellular structure and, after about 30 min tumbling movement, squeezed into the cellular structure. These data dynamically indicate that fusiform structures could directly transform into cellular structures. The size of this fusiform structure-derived cellular structure at 48 min was larger than that at 0 min. Also, other time-lapse images (lower panels, S. Fig. 4) showed two fusiform-connected cellular structures (thick arrows). The upper one connected with two fusiform structures (5 min, tick- and wide-arrows) both of that later squeezed inside this cellular structure. Again, the amplified images showed that the size of this newly formed cellular structure at 17 min was larger than that at 0 min (S. Fig. 4). These data further provide evidence that multiple fusiform structures participated in the formation of a cellular structure. Thus, we believe that mesenchymal cell progenitors originate from the transformation of fusiform structures.

**Figure 6.**
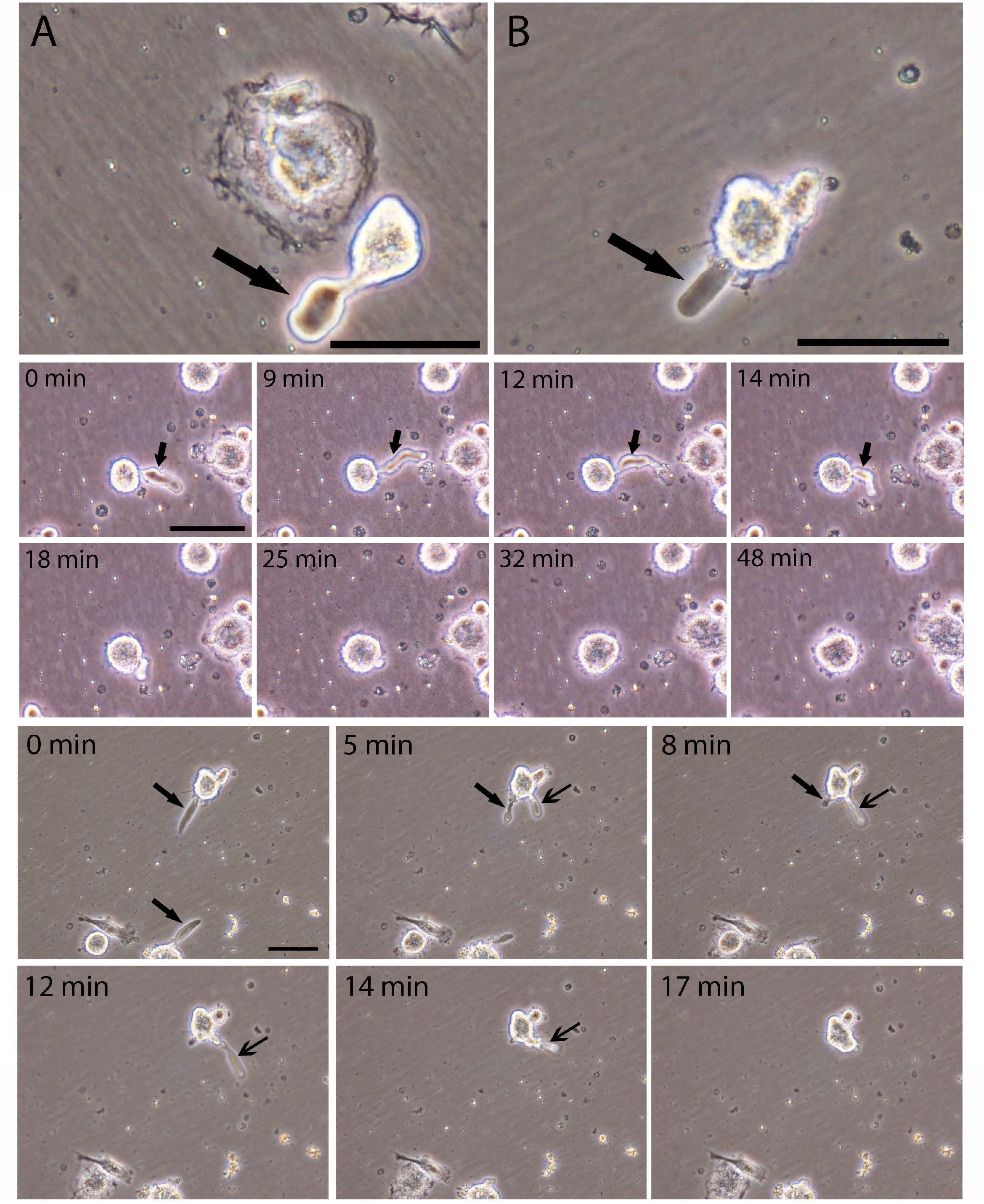
Cultured hUCB large fusiform structures directly transformed into mesenchymal cell progenitors. hUBC cells were cultured for 1 week and then observed under the microscope. Many large fusiform structures connected to the cellular structures (arrows in A, B) were identified. The middle panels showed the time-lapse images that a large fusiform structure (arrows) connected to a cellular structure and, after about 30 min tumbling movement, squeezed into the cellular structure. The lower panels showed two fusiform-connected cellular structures at time 0 (thick arrows). The upper cellular structure is connected with two fusiform structures at 5 min interval (tick- and wide-arrows). By the 17 min interval, both fusiform structures were inside this cellular structure. Bars in A, B, 0 min = 50 μm.

### Nucleated cells are required in the formation and differentiation of the mesenchymal cell progenitors

In our previous hUCB report (Kong et al., 2013a), we described how the cellular structures were formed by the fusion of multiple small particles, or NPRCPs. Also, we described that two or three medium-sized fusiform structures or PFDNCs were fused together or engulfed by each other to become one cellular structure. However, all the activities or the formation of the cellular structures needed the presence of eukaryotic cells or nucleated cells. After re-examining the previous hUCB video records, the time-lapse images (Fig. 7A, S. Fig. 5) showed the separated steps of how the new cells were formed in the presence of eukaryotic cells. Six medium-sized fusiform structures were numbered from #1 to #6, respectively; and a large fusiform structure-derived cellular structure was identified (wide-head arrow). All of them were located either on top or near a nucleated cell. Two fusiform structures (#1, and #2) were fused together to form a cellular structure by 59 min. The #6 turned into a small cellular structure by 9 min. The #3 was interacting with a group of particles during the time periods. The #4 and #5 were moving in and out of the nucleated cell, mostly around the nuclear area, (similar moving in and out of the nucleated cell activities were also seen inside the nucleated cell in Movie 1). The newly formed cellular structure (wide-head arrow) acquired the cell membrane and became a cell with a membrane by 92 min. These images clearly demonstrate that the medium-sized fusiform structures can increase their volume by collecting circulating particles (#3), acquiring materials from nucleated cells (#4, #5), fusing with each other (#1, #2), and finally acquire cell membrane from the eukaryotic cells (wide-head arrow). The medium-sized fusiform structures showed a quick and tumble-like movement (Movie 2), which made them easily approach the target cells to collect needed materials. Thus, due to the need for the materials from the nucleated cells, grouped fusiform structures were located mostly near or on top of a nucleated cell, as shown in Fig. 7B and 7C. Although the aggregated morphology was similar to that of the cell colonies, only one nucleus (thin arrow) could be observed in each group. We believe that these morphologies could be easily mistaken for stem cell colonies.

**Figure 7.**
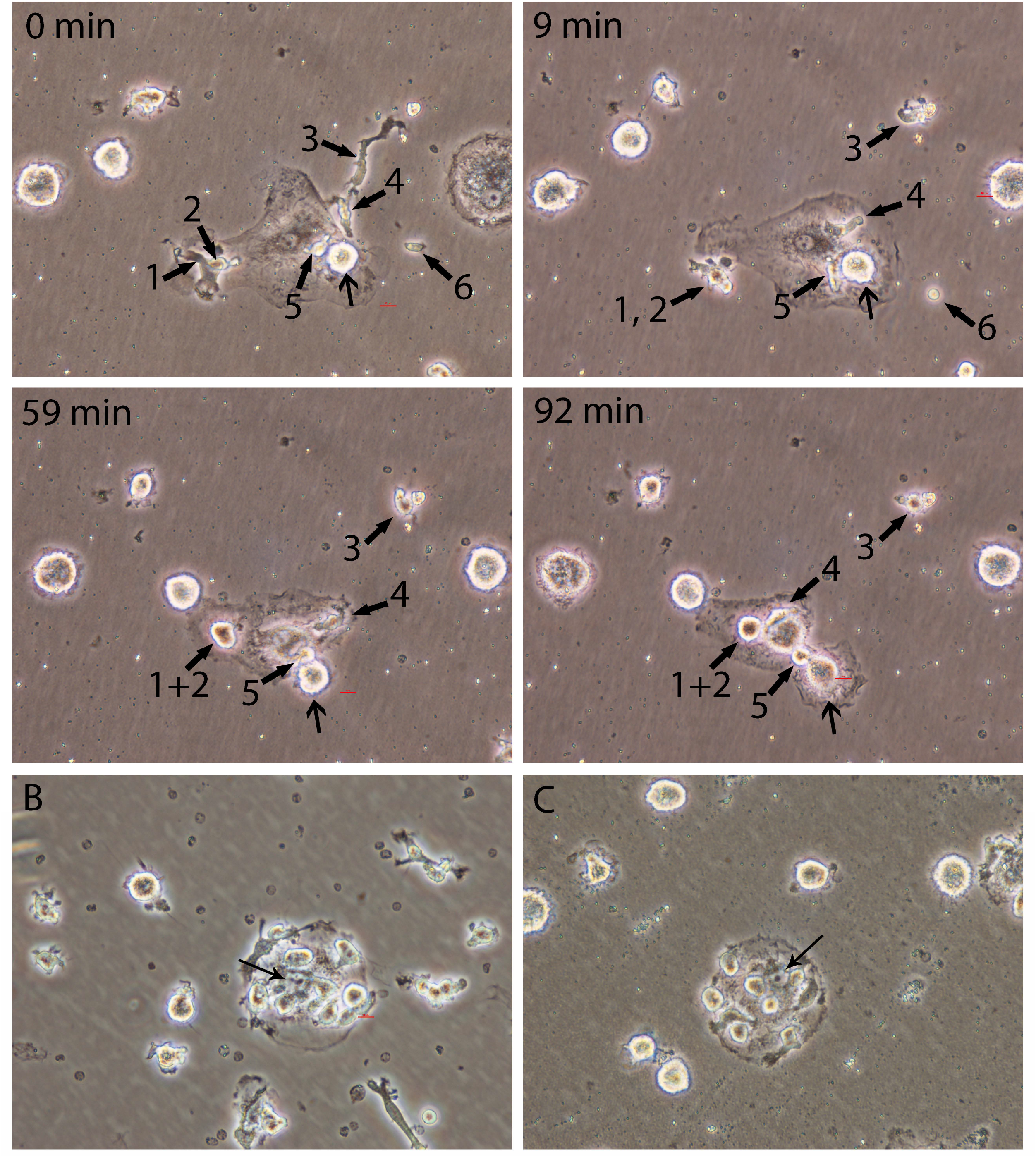
Nucleated cells are required in the formation and differentiation of the mesenchymal cell progenitors. The upper panels showed the time-lapse images of six medium-sized fusiform structures that were numbered from #1 to #6, respectively; and a large fusiform structure-derived cellular structure (wide-head arrow). Grouped fusiform structures were located mostly near or on top of a nucleated cell (B, C). Only one nucleus (thin arrow in B, C) could be observed in each group. Images were taken by a 40 x microscope lens.

### Nucleus formation after the cellular structures formed

The images in Fig. 2E – 2G, showed that the fusiform structures released from the semitransparent granules were in grouped patterns, and thus were similar to the pattern of stem cell colonies if cultural. In cultured hUCB, images confirmed that the grouped fusiform structures (arrow in Fig. 8A), and the grouped newly formed small cellular structures located near a larger cellular structure (arrows in Fig. 8B - 8D) resembled the morphologies of stem cell colonies. The medium-sized fusiform structures quickly transformed into a small cellular structure (S. Fig. 6). However we observed that none of these grouped cellular structures had a standard nucleus, or that they were not from stem cell mitotic division. To further confirm our statement, these newly formed cellular structures were over-stained with H&E, and Hematoxyline-only to examine the nuclear materials. Cellular structures containing granule materials or with protrusions on the surface areas (arrows in Fig. 8E - 8P) were considered as newly formed. Images showed that the newly formed cellular structures did not have any visible nucleus (Fig. 8E - 8H). Instead, all the larger-sized cellular structures had a nucleolus (Fig. 8I - 8L) that was surrounded by weak-stained nuclear materials. Hematoxyline-only stain also revealed that only a small amount of nuclear materials were in the newly formed cellular structure (Fig. 8M, 8N, 8P), but a nucleolus appeared in a larger-sized cellular structure (Fig. 8O). These data further confirmed that initially, the newly formed cellular structures did not have a nucleus.

**Figure 8.**
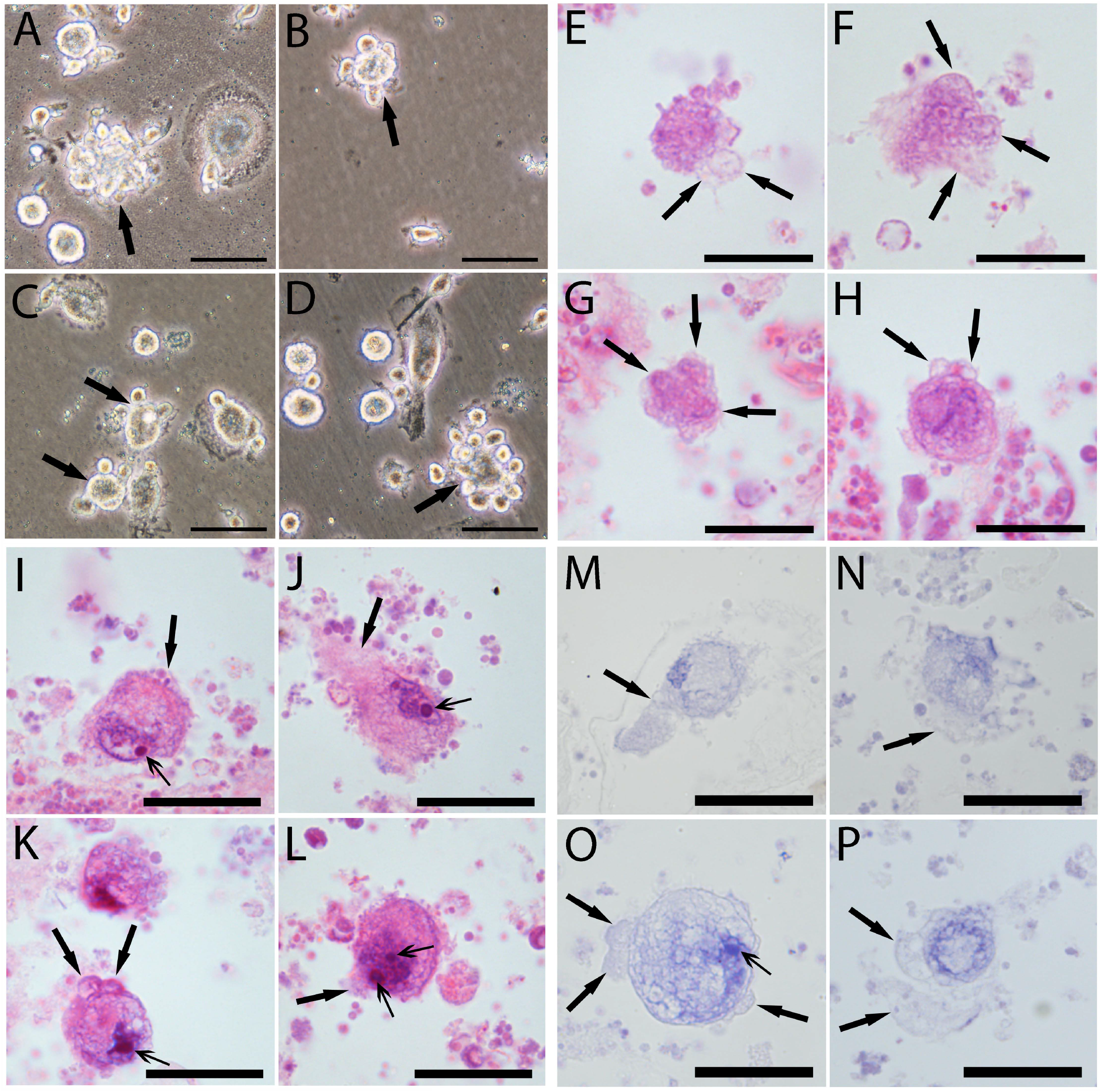
Nucleoli appeared before the nuclei in the new cellular structures. Grouped fusiform structures (arrow in A), and the grouped newly formed small cellular structures were identified in cultured hUCB (arrows in 8B-8D). Newly formed cellular structures were over-stained with H&E (E – L), and Hematoxyline-only (M – P) to examine the nuclear materials. Cellular structures with protrusions on the outer areas were considered the newly formed (thick arrows in E - P). The newly formed cellular structures did not have any visible nucleus (E - H). Instead, all the larger-sized cellular structures had one or two nucleoli (thin arrows in I - L). Hematoxyline-only revealed that only a small amount of nuclear materials were in the newly formed cellular structure (M, N, P), but a nucleolus appeared in a larger-sized cellular structure (thin arrow in O). Bars in A-D = 50 μm; in E-P = 20 μm.

### Human mesenchymal stem cells were from the belt-shaped areas

During hUCB culture, we often observed, after one week of culture, some long-belt-shaped areas that were covered with crowded small sand-like particles, fusiform structures, and large cellular structures (Fig. 9A, 9B, arrows in S. Fig. 7), which had a lot of similarities to the composition of the long segmented materials observed in mouse blood (Fig. 2). The belt-shaped areas were longer than 1 mm (S. Fig. 7A, 7B), had more cellular structures on its top than in other areas. Although some fusiform structures still could be observed (arrow in Fig. 9A) after one week in culture, the majority of populations on the top of the belt-shape area were the newly formed large and small cellular structures (Fig. 9A, 9B). Time-lapse images showed that small cellular structures were constantly migrated out of the belt-shaped area (low panel in Fig. 9, S. Fig. 8), suggesting that they originated from the belt-shaped area. We believe that the components in the belt-like patterns were the same as the long segmented materials that could further develop after attaching to the culture plates.

**Figure 9.**
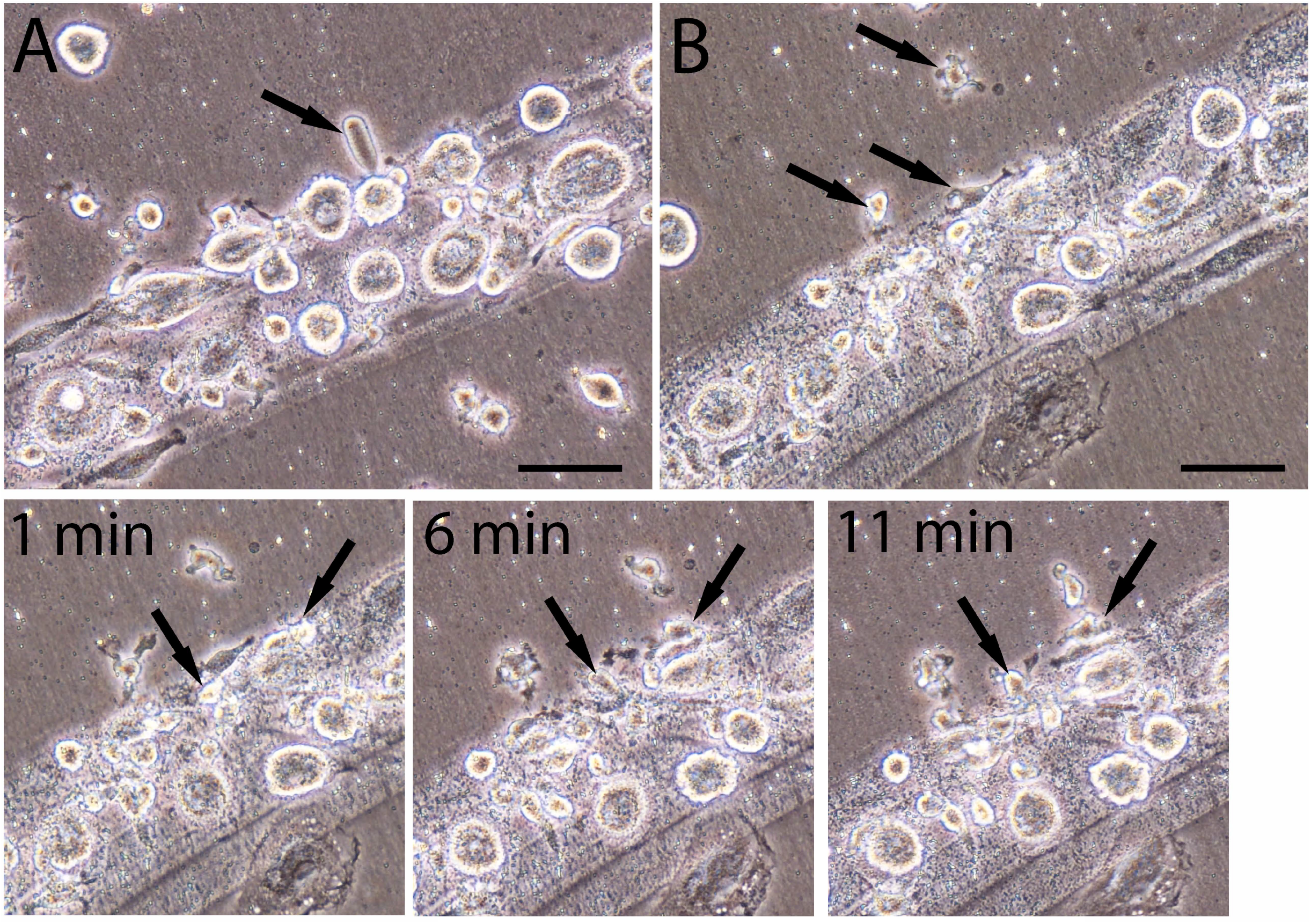
Human mesenchymal stem cells were also from the long segmented materials. Long-belt-shaped areas were observed in one-week cultured hUCB. These areas were covered with crowded small sand-like particles, fusiform structures (arrows in A, B), and large cellular structures. Time-lapse images showed that small cellular structures were constantly migrated out of the belt-shaped area (low panel). Bars in A, B = 50 μm.

### The black segments were released from the tube-shaped structures

The identification of the long segmented materials in mouse blood and the belt-like patterns in the cultured hUCB suggested that they were from long-shaped origins. Thus, we asked if the long segmented materials were from the tube-shaped structures similar to that in our previous report (Kong et al., 2019). We then, using H&E stains in mouse blood, identified a few dark blue-colored tube-shaped structures each of that had a fully spiral-shaped opening, suggesting that it had released its inclusions (Fig. 10A, S. Fig. 9). High magnification images still identified traces of black sand-like materials in the opening crack (arrow in 10C) or on top of the tube-shaped structure (arrows in 10B, 10D). Thus, we believe that the dark blue-colored tube-shaped structure was the origin of the dark segments or was the niche of the mesenchymal stem cells. The dark blue-colored tube-shaped structures were more than 1 mm in length, suggesting that one of the tube-shaped structures could release numerous fusiform structures and eventually produce thousands of mesenchymal cells.

**Figure 10.**
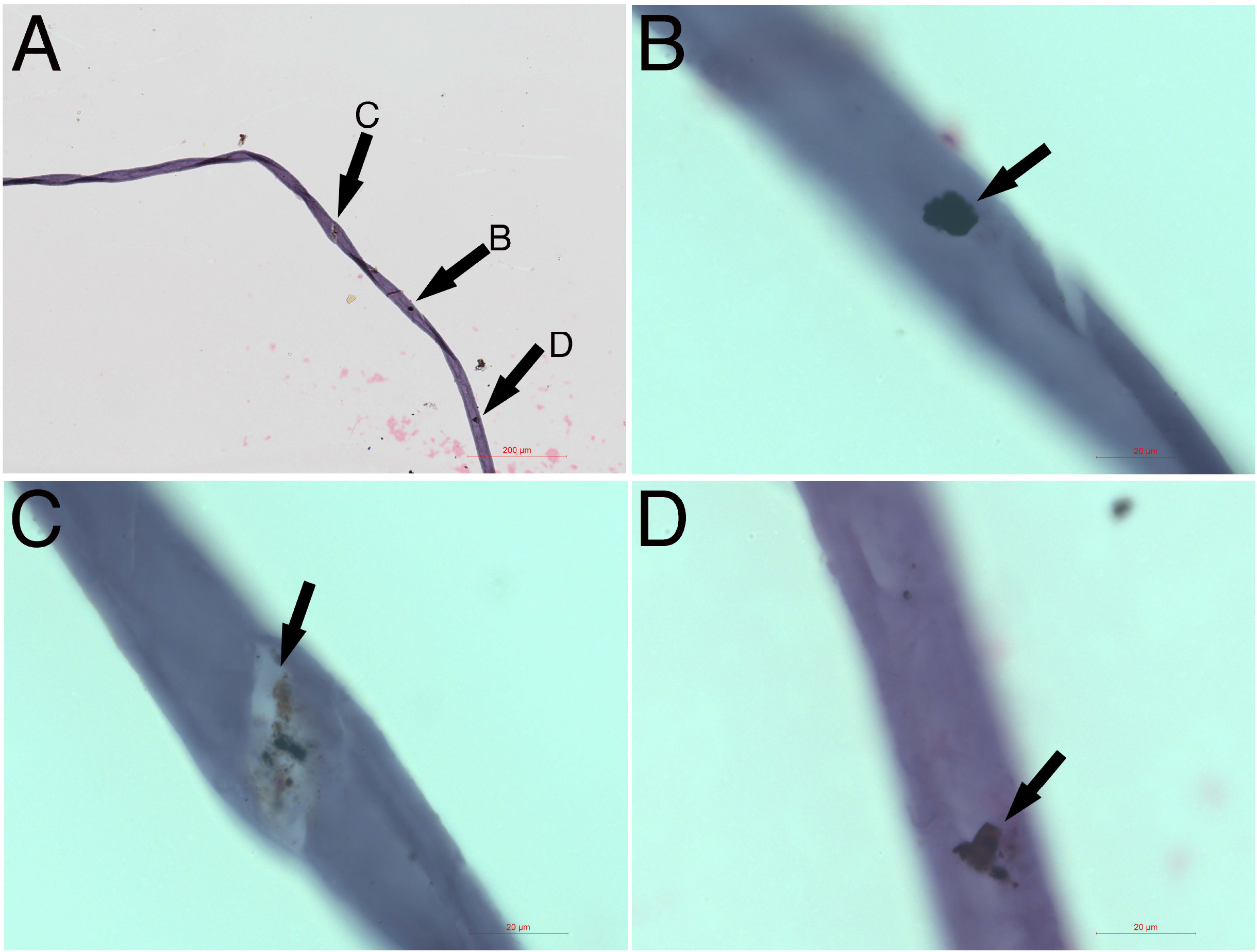
The black colored materials were from the tube-shaped niche. A dark-blue colored tube-shaped structure that was more than 1 mm in length and had a spiral opening was identified (A). High magnification images identified traces of black debris in the opening crack (arrow in C) or on top of the tube-shaped structure (arrows in B, D). Bar in A = 200 μm; in B-D = 20 μm.

## Discussion

Our data provide evidence that the origin or the niches of mesenchymal stem cells are tube-shaped tissues. The contents inside the niches need at least 5 procedures to develop into the new mesenchymal stem cells. These procedures include the release of the segmented materials, release of the fusiform structures, fusion of fusiform structures, transformation of the cellular structures, and nucleus formation, however, not by mitosis. Our report revealed these step-by-step procedures of how the new mesenchymal cells are formed. In addition, the segmented materials and the fusiform structures are first observed and described in this report. Thus, we believe that our reports would contribute new information in the stem cell field.

Similar tube-shaped structures have been reported in our previous study (Kong et al., 2019). We have reported two morphologically distinct tube-shaped structures, light purple color, and copper color, and their released materials are termed progenitor-releasing filaments and bud-shaped structures. These structures eventually produce lineage-specific CD34-positive stem cells. Here, we show the dark blue-colored tube-shaped structures that produce and release the segmented materials that eventually form into mesenchymal stem cells. Thus, we believe that the tube-shaped structures are the origins, or the niches, of all or most lineage-restricted stem cells. The location of these tube-shaped MSC niches is still unclear. But, we question the dogma that the niche of MSC locates in the bone marrow because we did not find bone marrow is rich in the dark fragmented materials or the dark blue-colored tube-shaped MSC niches. The tightly enclosed long tube-like structure suggests that any cells or factors inside the lumen are completely isolated from outer environments. We believe that one type of tube-shaped structure produces only one type of lineage-specific stem cells, or that the tube-shaped structures are lineage-restricted. Although until now, we can only identify or distinguish them based on their distinct color of H&E stains, the size of the width, and the length. We believe that at least two hundred types of tube-shaped structures are present in the human body because there are at least two hundred cell types in human body. Also, each type of lineage-specific tube-shaped structure must be rich in its specific tissues.

Identification of the fusiform structures is a novel finding. We have observed that only the dark segmented materials release the fusiform structures, or that the fusiform structures are found only in the process of mesenchymal cell formation. The larger fusiform structures can directly transform into cellular structures and the small ones can fuse together to further transform into cellular structures. These cellular structures eventually develop into mesenchymal stem cells. In our previous study (Kong et al., 2013a), we described how the new Oct4 and Sox-2-positive stem cells are formed by the fusion of particles, the NPRCPs, in hUCB. Now we believe that NPRCPs are small fusiform structures, which is hard to distinguish from the circulating particles. The mechanism of particle fusion to form new kidney epithelial cells and endothelial cells has been described in our previous studies (Kong et al., 2013b; Kong et al., 2017). However, we still question if the mechanism of particle fusion in the mesenchymal stem cell formation is the same mechanism to form other specific stem cells, because the fusion of particles to form into these new stem cells appeared via distinct different procedures.

Due to the sizes of the small fusiform structures, they could be easily considered as particles. Thus, this report described three particle-like materials, the small-sized fusiform structures that were flouting in the lined groups (Fig. 2B, 2H), the sand-like particles in the fragmented materials or the belt-shaped areas (Fig. 2A, Fig. 9), and the particles inside the large-sized cellular structure (Fig. 4F). Although all these three particles may have biological activities, only the small-sized fusiform structures can participate the fusion and further become the contents in the new cellular structures. The function of sand-like particles in the segmented materials is still unclear but possibly involved in providing the biological materials for, or participating in the development of the new cells and the differentiated cells. The particles that popped out of the newly formed cellular structures are possibly from the fused small fusiform structures or collected circulation particles. We believe that these particles inside the newly formed cellular structures only participate in the development of the new cells. Because the newly formed cellular structures are not eukaryotic cells, thus do not possess the intracellular organelles or the intracellular pathways to release biological materials, or they are only taking, not giving. In recent years, it has been well claimed that mesenchymal stem cells release exosomes that have positive effects on tissue regeneration or cell therapy studies. However, we have observed that most circulation particles are either in the process of forming the new cells by direct transformation or by fusion to participate in the formation of new stem cells. The direct transformation by the 1 um diameter “spore-like” pre-cells have been reported in our previous report (Kong et al., 2019). In comparison, it is rare to observe numerous particles released from the eukaryotic cells. Thus, we question the theory of the exosomes released by mesenchymal stem cells. We believe that the circulation particles are a mixed complicated population and need to be clearly sorted out in future studies.

Our observations strongly suggest that the new mesenchymal stem cells are from the fusion of multiple small fusiform structures and the transformation of large fusiform structures. However, the mechanism of the nucleus formation in the cellular structures is unclear. We have observed that the nucleolus is the main nuclear material in the early stages of the newly formed cellular structures, or the nucleolus appears before the nucleus appears. Nucleoli contain DNA, RNA, and proteins, and are best known as the sites of ribosome biogenesis (O’Sullivan et al., 2013). We hypothesize that the nucleolus observed in the newly formed cellular structures is possibly the first-made intracellular organelle. Although until now we still don’t know how the nucleolus gene was obtained in the newly formed cellular structures in the present study. We speculate that the DNA fragments in the apoptotic cells may be the origin of the nucleolus in the new cells because we have observed that the DNA in the small “spore-like” pre-cells was from apoptotic cell DNA fragments (Kong et al., 2019). Recently, some reports indicate that apoptotic cells are needed for the therapeutic effects of MSC (Pang et al., 2021). Although their explanation for the function of the apoptotic cells is not the same as ours, it is positive that apoptotic cells, in some way, participate the regenerative function of MSCs, which further supports our hypothesis that the lineage-specific apoptotic cells that contain DNA fragments, participate the formation of the lineage restrict tube-shaped niches and provide their DNA fragments for the nucleus formation in the newly formed lineage-restricted stem cells.

In the past decades, the therapeutic effects of MSC have been credited from the cells to the MSC-released exosomes. Our findings clearly described the renewal procedures of MSCs, which could explain why the attention to MSCs has been shifted to exosomes. We believe that the newly formed cellular structures or the new cells containing only nucleolus do not have a standard cell membrane, and thus could not be recognized and rejected by the recipients. Our data also show that circulation contains many small lineage-specific particles or pre-cells; thus, their therapeutic effects should not be credited to the multifunction of MSCs or exosomes. Again, our data indicate that newly formed MSCs are not from the mitotic division.

## Abbreviation

MSCs: (Mesenchymal stem cells)
hUCB: (human umbilical cord blood)
PBS: (phosphate buffered saline)
H&E stain: (Hematoxylin and eosin stain)
NPRCPs: (Non-platelet RNA-containing particles)
PFDNCs: (particle fusion-derived non-nucleated cells)

## Declarations

### Ethics approval and consent to participate

Not applicable.

### Consent for publication

Not applicable.

### Availability of data and materials

All data generated or analyzed during this study are included in this published article (and its supplementary information files).

### Author Contributions

W. Kong: Manuscript writing, Project directing, Experimental designs.

X. Han: Performing experiments, Data collection.

H. Wang: Performing experiments, Data collection.

X. Zhu: Performing experiments, Data collection.

## Supplementary data

**S. Fig. 1. Identification of segmented materials.** Dark-colored segmented materials were identified with H&E stains in mouse blood. The sizes of these segments, although differ in length, but similar in width (A – D). Bars = 50 μm.

**S. Fig. 2. Light-weighted fusiform structures arranged in a line pattern.** A group of fusiform structures arranged in a line pattern was identified on the top of the tissue slides. The pink-stained shadows in the background suggested that the stained cells were not at the same levels as the fusiform structures. Bar = 50 μm.

**S. Fig. 3. A large fusiform structure was connected to a round thin membrane that contained sand-like particles.** Bar = 50 μm.

**S. Fig. 4. Multiple fusiform structures can transform into one larger cellular structure.** Detailed time-lapse images for the lower panel images are in Figure 6. The amplified images for the low panels in Fig. 6 showed that the size of this newly formed cellular structure at 17 min was larger than that at 0 min. Bars = 50 μm.

**S. Fig. 5. The detailed time-lapse imaged to support the upper panel images in Figure 7.**

**S. Fig. 6. Medium-sized fusiform structures transform into the small cellular structure.** Time-lapse images showed that the medium-sized fusiform structures transformed into a small cellular structure in 29 min. Bar = 50 μm.

**S. Fig. 7. Belt-shaped areas were identified in the cultured hUCB.** In the cultured hUCB, the belt-shaped areas were identified (A, B). These areas are covered with crowded small sand-like particles (arrows), fusiform structures, and large cellular structures (C, D). Bars in A and B = 200 μm; in C and D = 50 μm.

**S. Fig. 8. Human mesenchymal stem cells migrated out of the belt-shaped areas**. The detailed time-lapse images for the images in the low panel in Fig. 9 showed how the fusiform structures or newly formed cellular structures migrated out of the belt-shaped areas.

**S. Fig. 9. The blue-colored tube-shaped structures were the origin of the segmented materials.** A spiral-shaped open tube-shaped structure that stained dark blue with H&E, about 600 μm was identified in mouse blood. Bar = 200 μm.

## Movie Legends

Movie 1. The video showed that a fusiform structure was directly connected, not stuck on, to a cellular structure. The cellular structure had numerous small particles in the center. The video was taken by a Nikon inverted microscopy with 40x lens and zoomed 4 times.

Movie 2. In cultured hUCB, the medium-sized fusiform structures were moving extremely active comparing to the cells in the same area. The video was taken by a Nikon inverted microscopy with a 40x lens.

