## Supplementary Figures for "The self-renewal procedures of mesenchymal stem cells in the blood"

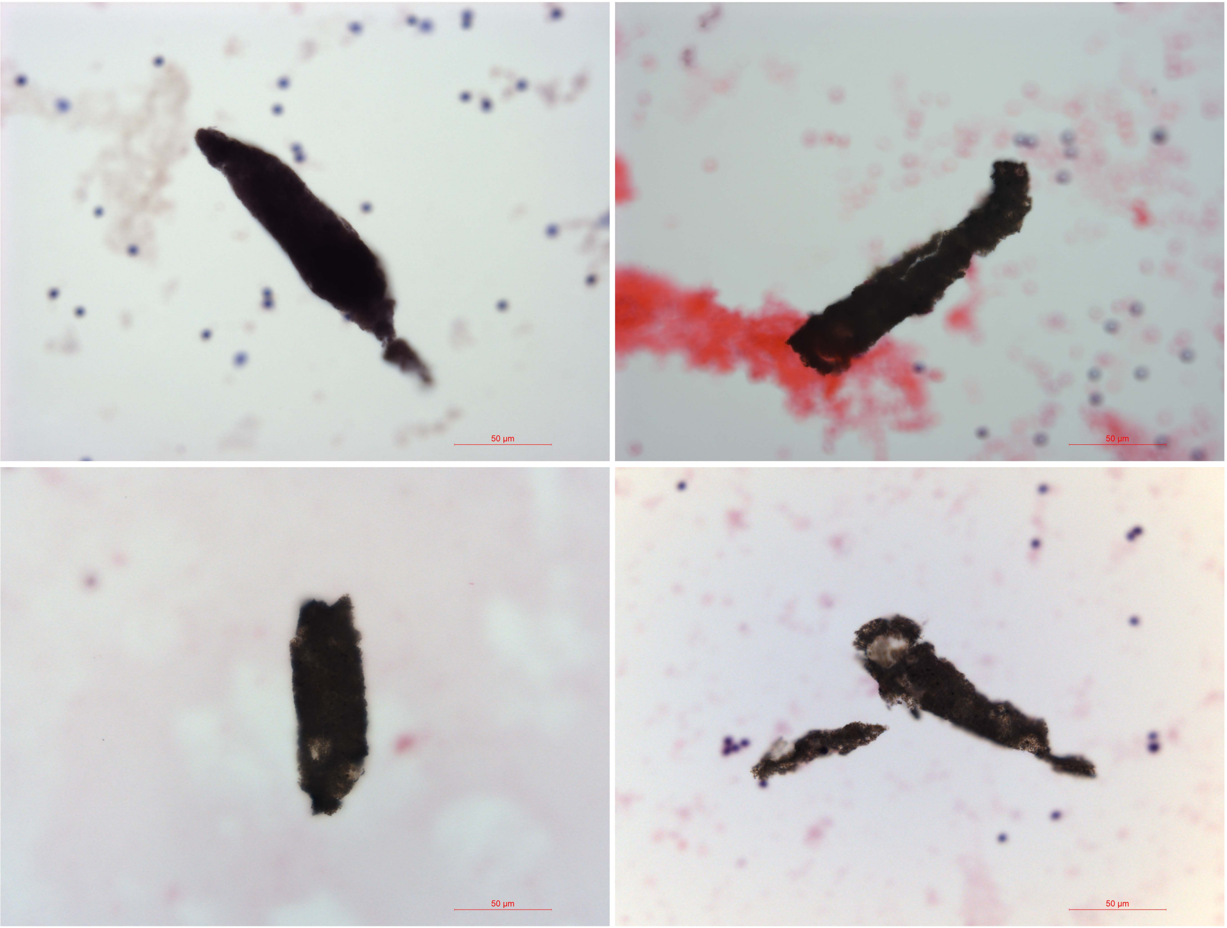


Supplementary Figure 1

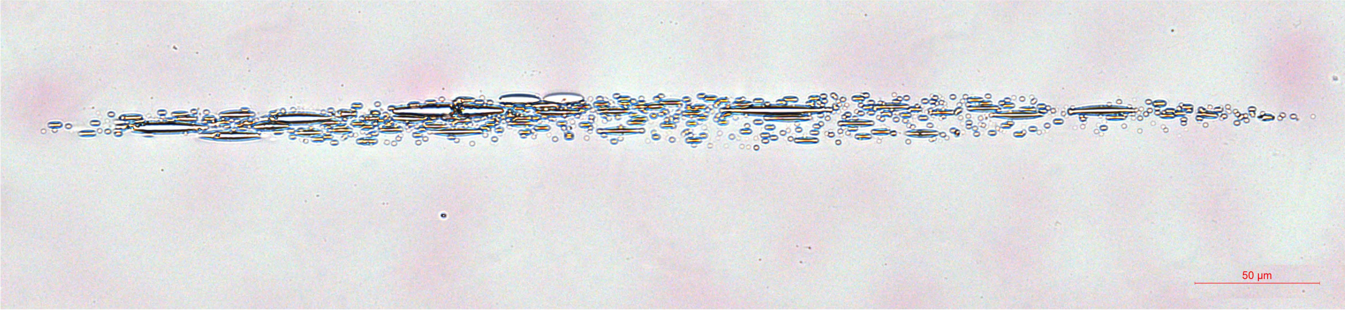


Supplementary Figure 2

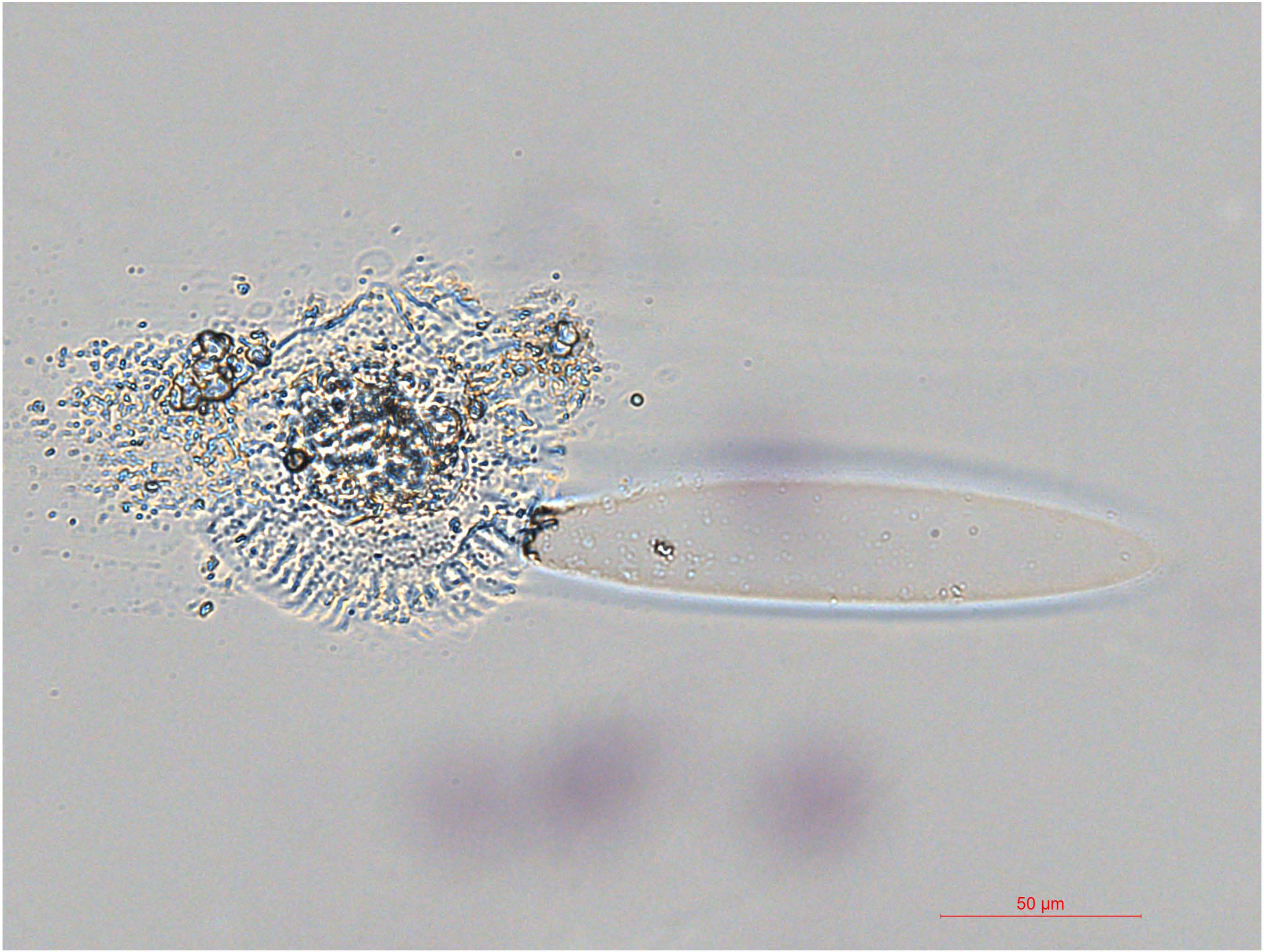


Supplemetary Figure 3

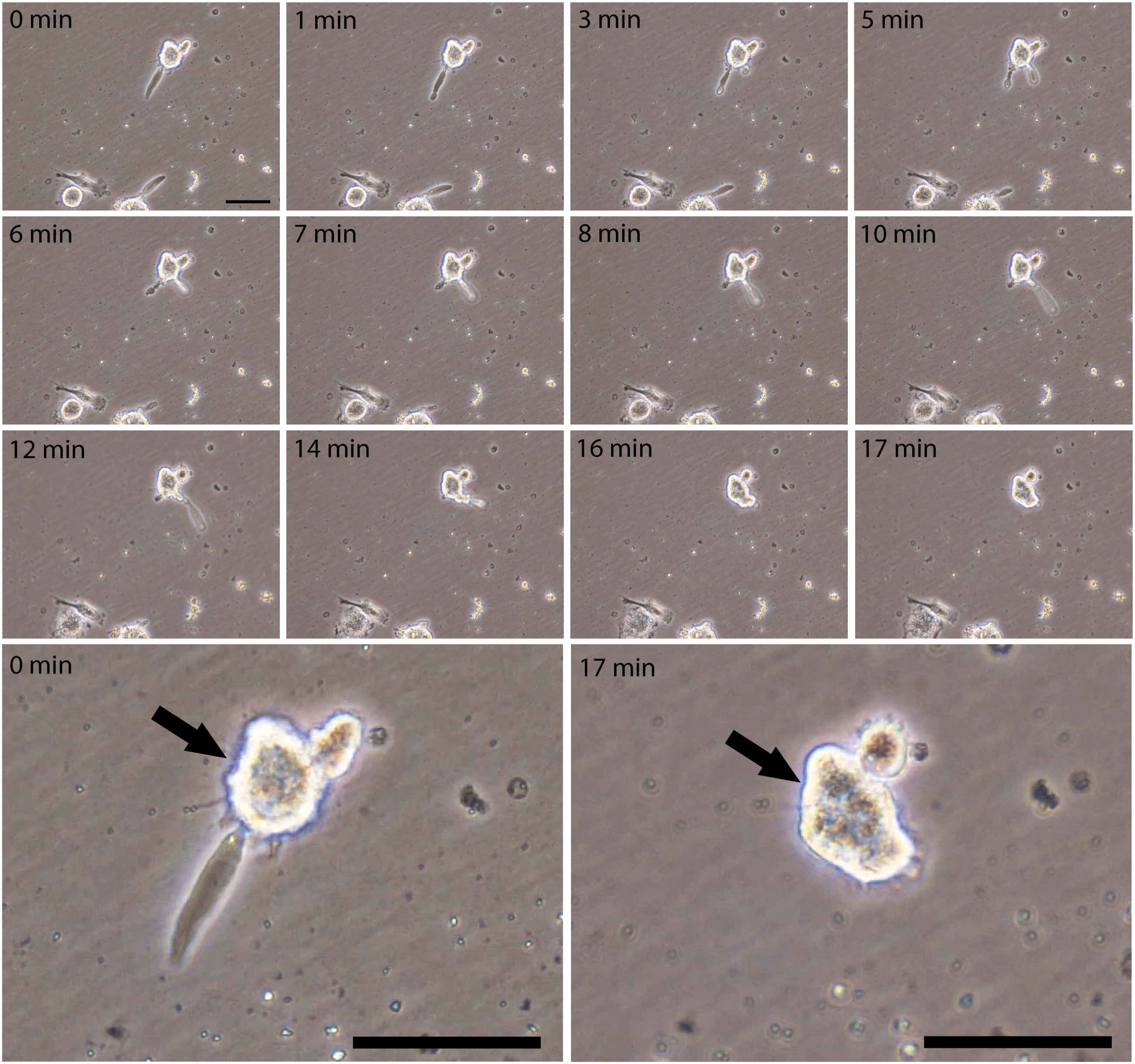


Supplementary Fig. 4

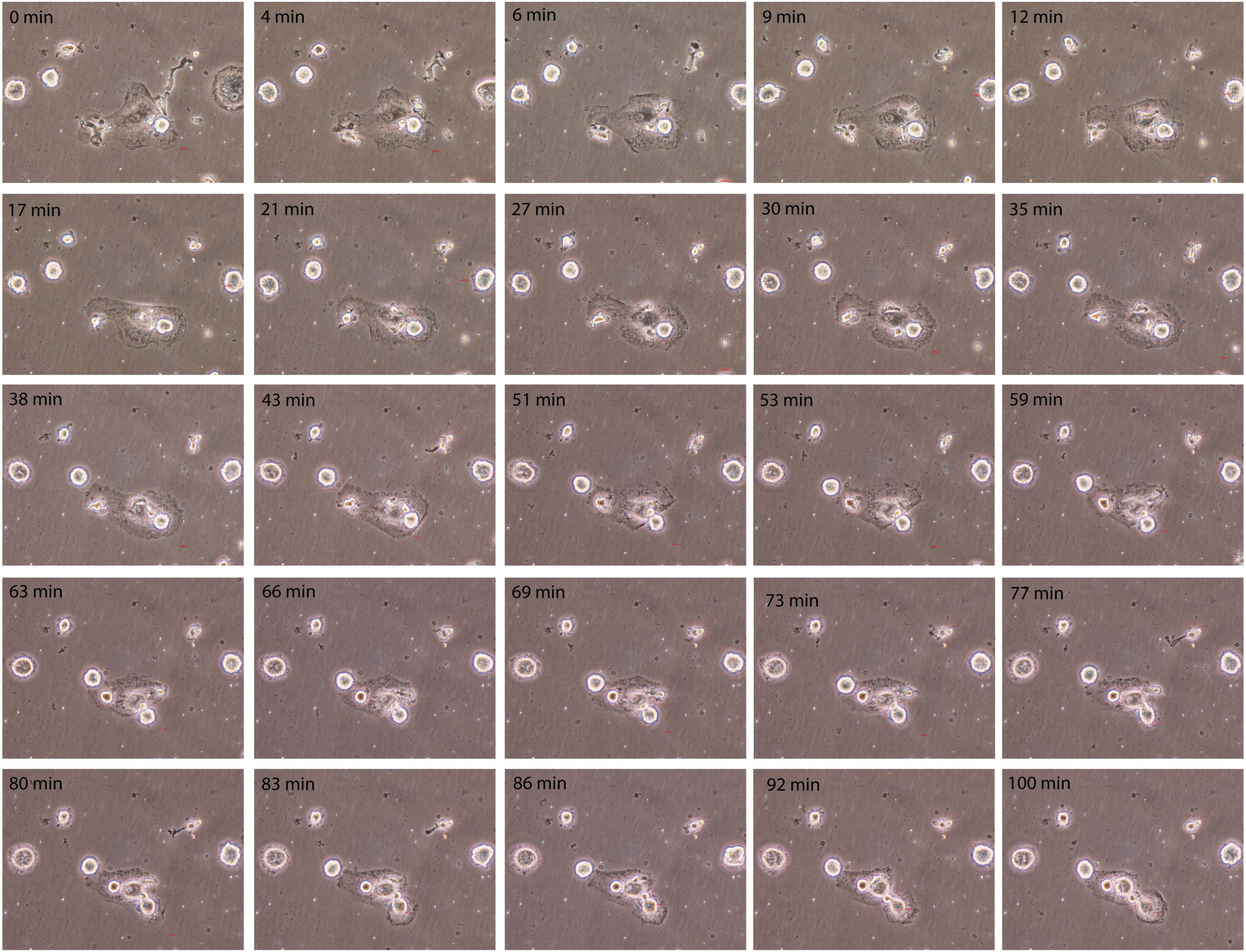


Supplementary Fig. 5

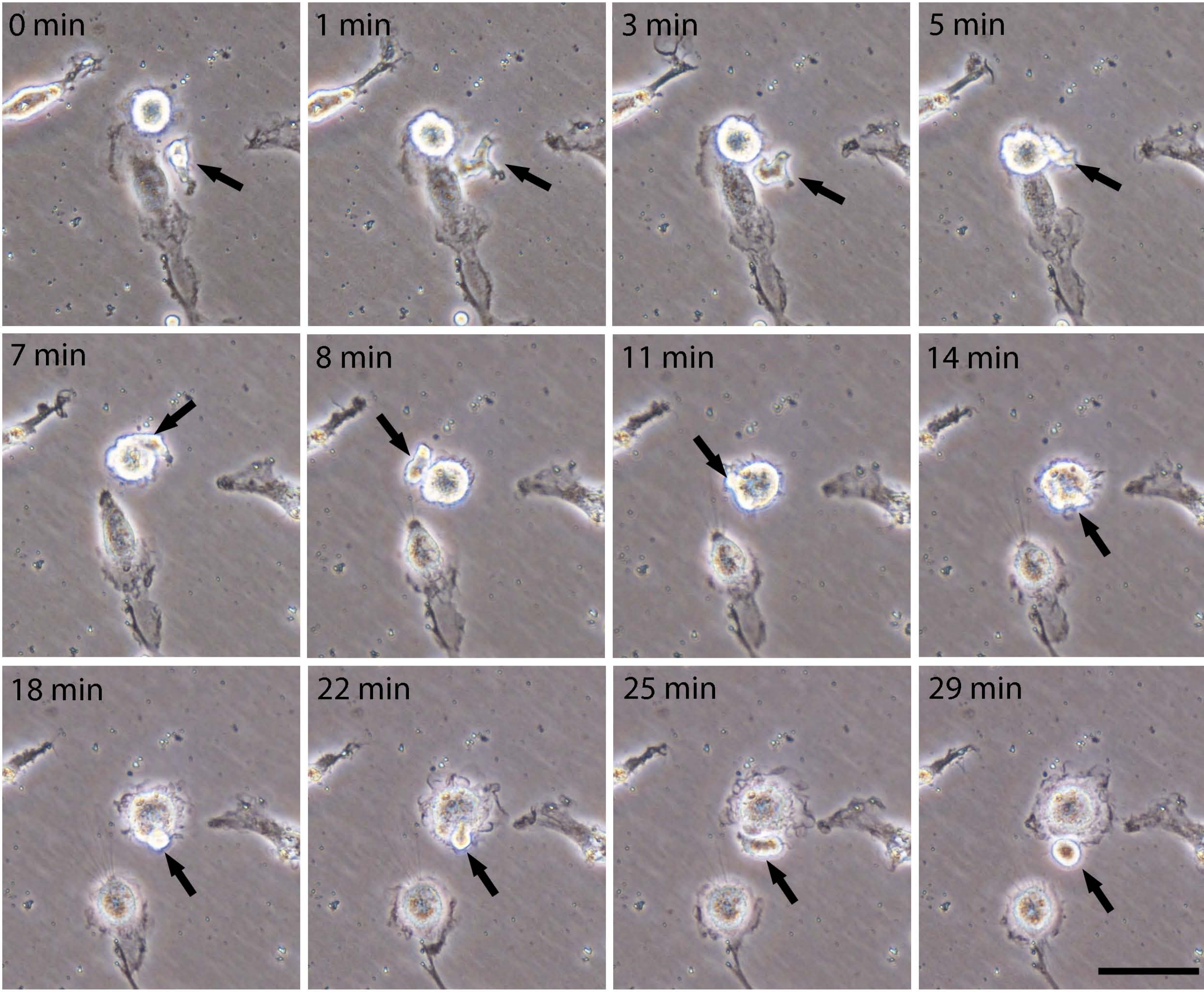


Supplementary Figure 6

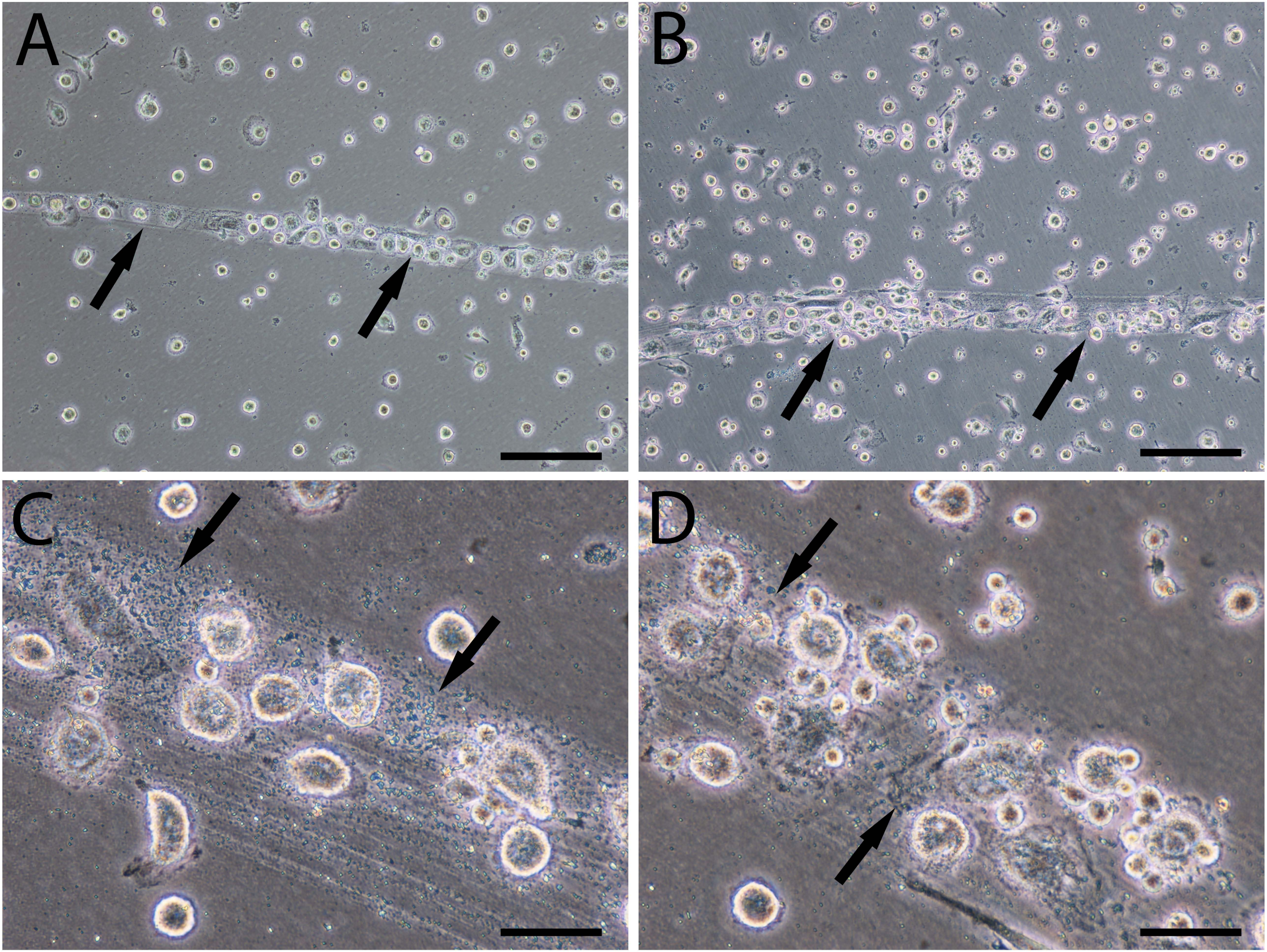


Supplementary Figure 7

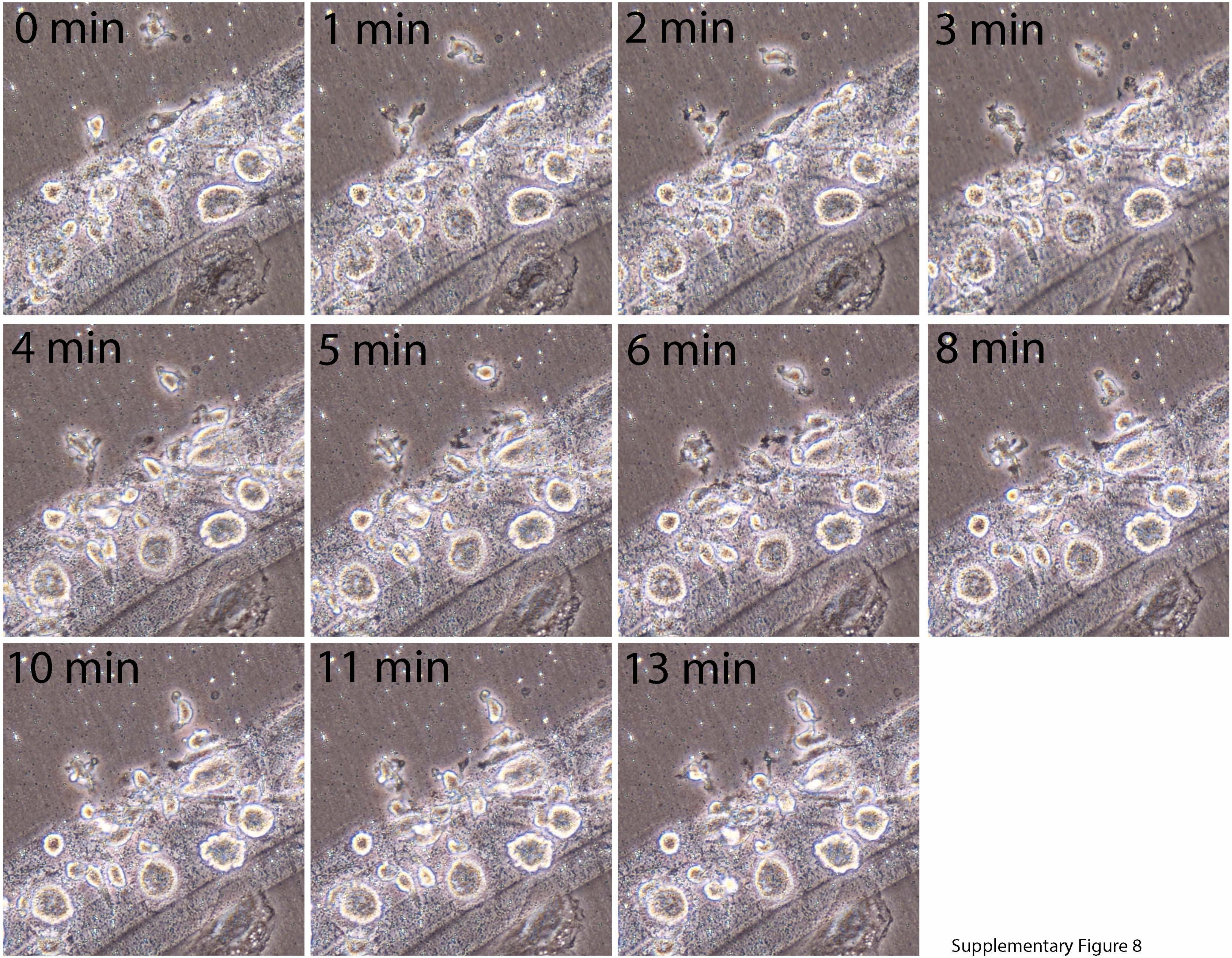


Supplementary Figure 8

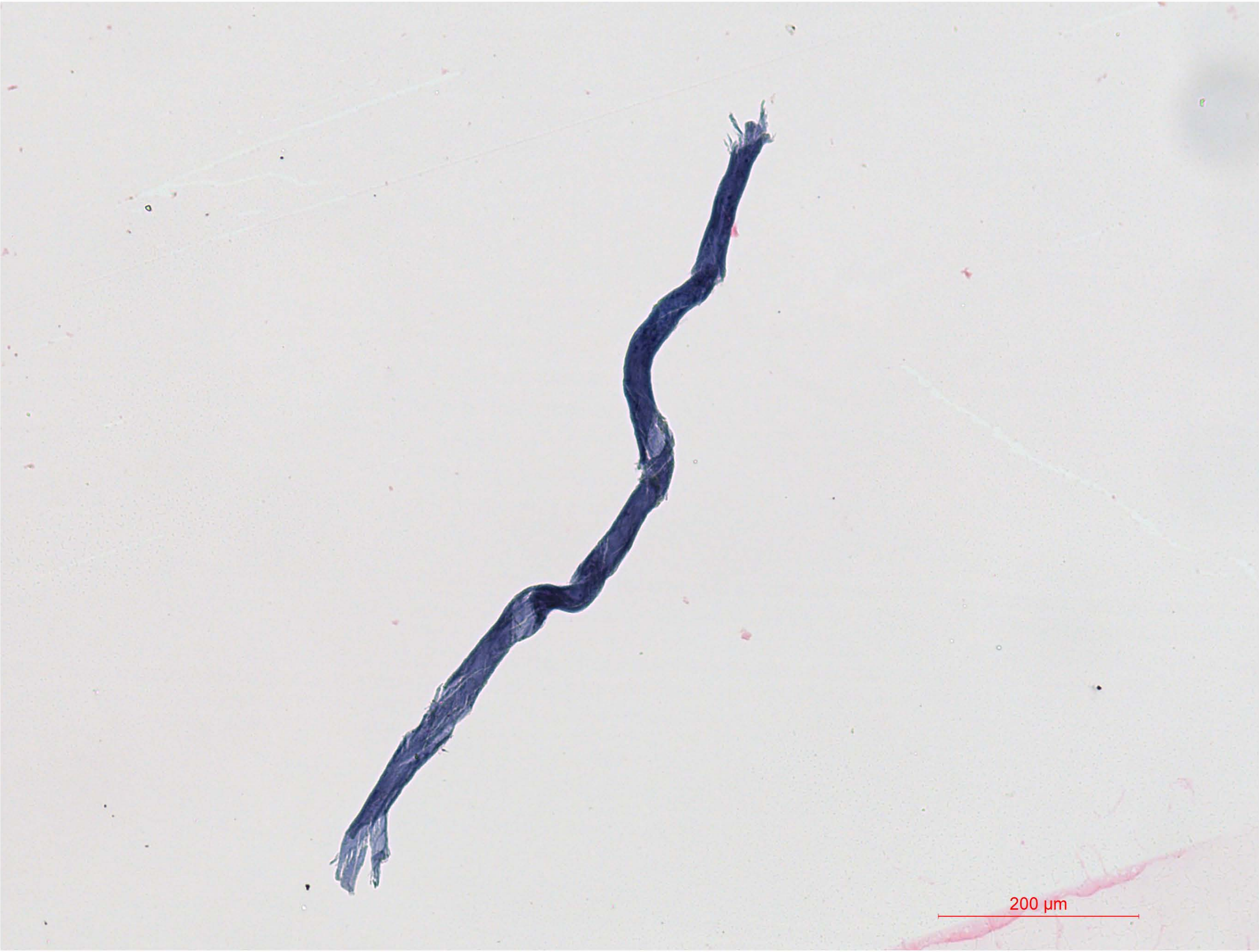


Supplementary Figure 9
